# Protein–Ligand Affinity Prediction via Jensen–Shannon Divergence of Molecular Dynamics Simulation Trajectories

**DOI:** 10.1101/2025.02.17.638772

**Authors:** Kodai Igarashi, Masahito Ohue

## Abstract

Predicting the binding affinity between proteins and ligands is a critical task in drug discovery. Although various computational methods have been proposed to estimate ligand target affinity, the method of Yasuda et al. (2022) ranks affinities based on the dynamic behavior obtained from molecular dynamics (MD) simulations without requiring structural similarity among ligand substituents. Thus, its applicability is broader than that of relative binding free energy calculations. However, their approach suffers from high computational costs due to the extensive simulation time and the deep learning computations needed for each ligand pair. Moreover, in the absence of experimental Δ*G* values (oracle), the sign of the correlation can be misinterpreted. In this study, we present improvements to Yasuda et al.’s method. Our contributions are threefold: (1) By introducing the Jensen–Shannon (JS) divergence, we eliminate the need for deep learning-based similarity estimation, thereby significantly reducing computation time; (2) We demonstrate that production run simulation times can be halved while maintaining comparable accuracy; and (3) We propose a method to predict the sign of the correlation between the first principal component (PC1) and Δ*G* by using coarse Δ*G* estimations obtained via AutoDock Vina.

## 1 Introduction

Protein–ligand binding affinity prediction is a cornerstone technique in drug design and biological research. Binding affinity information is essential for screening candidate compounds that either bind to a specific protein or target a specific ligand. However, large-scale screening using biochemical experiments or structural analyses requires significant time and effort. Thus, computational approaches for rapid and efficient screening are highly desirable [1].

Existing computational methods for binding affinity prediction can be broadly classified into physics-based approaches and data-driven approaches. Physics-based methods, such as absolute free energy calculations [2], free energy perturbation (FEP) methods [3, 4], and MM/PBSA [5], can yield highly accurate predictions but at a high computational cost. In contrast, data-driven approaches using machine learning or deep learning techniques [6–9] are typically faster during inference but require large, highquality training datasets and are often subject to biases present in the training data.

An alternative strategy evaluates binding affinity by comparing the dynamic behavior of proteins upon ligand binding using molecular dynamics (MD) simulations [10]. In this approach, MD simulations are performed for both the apo form of the protein and its various ligand-bound complexes. The differences in the dynamics of binding site residues between ligand systems are quantified using Wasserstein distances computed via deep learning [11]. The resulting distance matrix is embedded into a three-dimensional space using nonlinear dimensionality reduction, followed by principal component analysis (PCA). Finally, the value along the first principal component (PC1) is used to predict lower Δ*G* values. Yasuda et al. demonstrated good correlations between PC1 and experimental Δ*G* for targets such as Bromodomain 4 (BRD4) (*R* = 0.88) and protein phosphatase 1B (PTP1B) (*R* = 0.70). Nonetheless, the method is hampered by high computational costs from both the MD simulations and the deep learning computations required for each ligand pair. Moreover, even if PC1 values can be obtained, the sign of the correlation with Δ*G* remains ambiguous without oracle data.

In this study, we address these issues by reducing computational cost and expanding the method’s applicability. Specifically, we avoid the deep learning-based Wasserstein distance computation by employing the JS divergence for estimating similarity between trajectories. In addition, we propose a method to determine the sign of the correlation between PC1 and Δ*G* by using coarse Δ*G* estimations from docking simulations.

## 2 Materials and Methods

Figure 1 presents an overview of the proposed method. While the overall framework is similar to the method of Yasuda et al. [10], the key differences are in the computation of the distance matrix (step d) and the estimation of the sign of the PC1 correlation (step g). In our method, deep learning is replaced by the JS divergence for rapid evaluation of similarity between trajectories.

**Figure 1:**
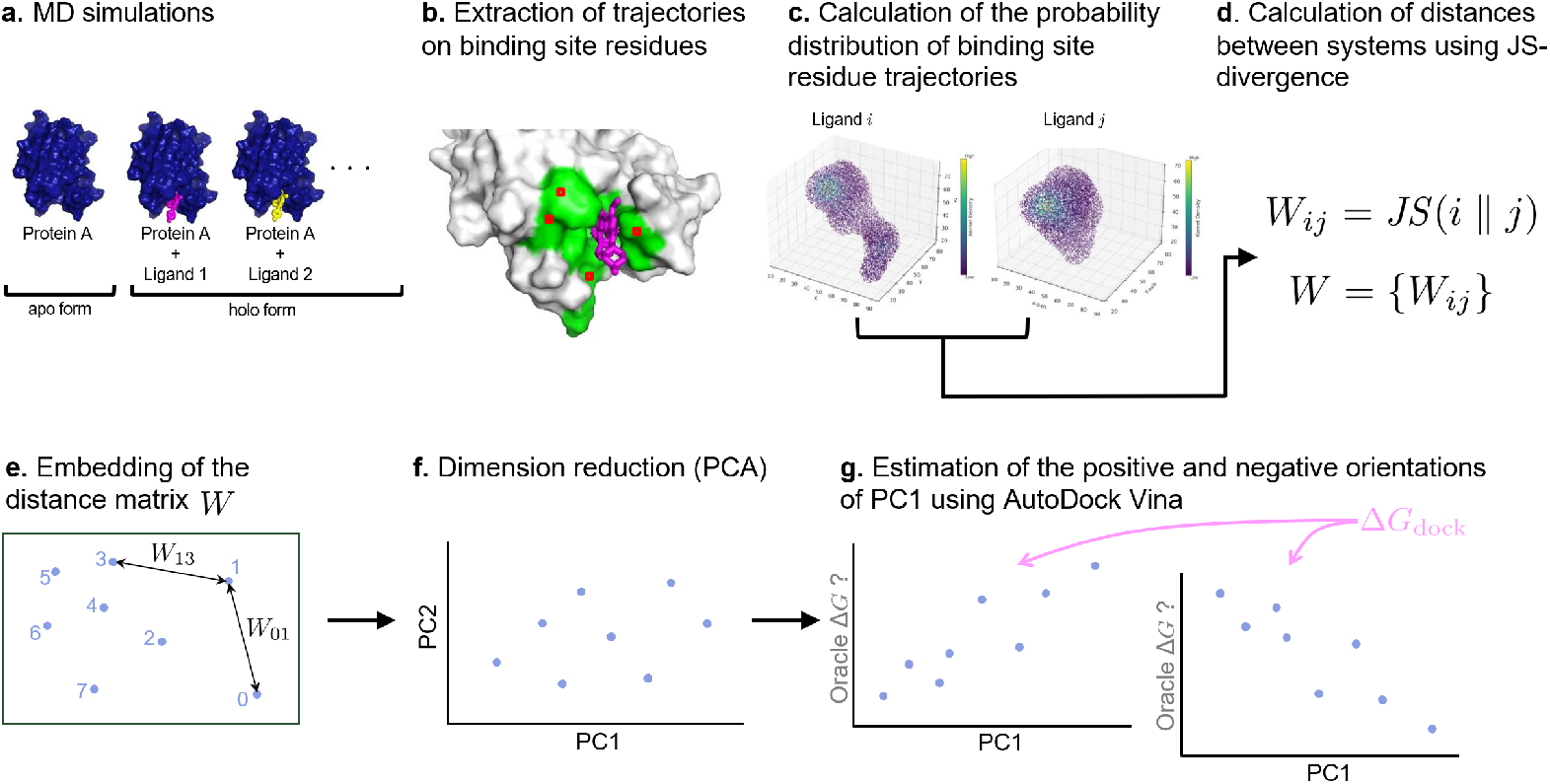
Overview of the proposed method. The main differences compared to the method of Yasuda et al. [10] are the computation of the distance matrix *W* (step d) and the estimation of the polarity of PC1 (step g).

### 2.1 Dataset

We evaluated the proposed method using three datasets: BRD4 and PTP1B as used in previous studies, and c-Jun N-terminal kinase (JNK1), which has been employed as a benchmark in FEP calculations [12]. These datasets are widely used in free energy calculations [2, 4] and are appropriate for evaluating the proposed method. The crystal structures of the complexes and the chemical structures of the compounds are provided in Supplementary Figures S1–S4.

### 2.2 Preparation of Initial Protein–Ligand Complex Structures

For BRD4, initial structures were generated from the crystal structures provided in the relevant PDB files [2]. For PTP1B and JNK1, the initial ligand positions were determined based on a reference protein structure [4]. Specifically, PTP1B used the crystal structure with PDB ID 2QBS, while JNK1 used the crystal structure with PDB ID 2GMX. Hydrogen atoms were added to each PDB file using H++ [13] at pH 7.0 to complete missing atoms.

### 2.3 Molecular Dynamics Simulation

All-atom MD simulations were conducted for both the protein and the protein–ligand complexes. Each hydrated system was subjected to energy minimization, followed by restrained MD simulations in the NVT and NPT ensembles, and finally a production run.

A cubic box with a 10.0 Å buffer region was created using the TIP3PBOX model [14] for solvation. The ff14SB and GAFF force fields were applied for the protein and ligand, respectively. The system charge was neutralized by adding sodium or chloride ions, and topology and coordinate files were generated.

Using Amber22 [15], each system was first minimized using 5000 steps of steepest descent. Subsequently, a 100 ps NVT simulation at 300 K was performed with a restraint of 10 kcal/mol ·Å^2^ on heavy atoms. This was followed by a 100 ps NPT simulation under the same restraint at 300 K and 1 bar.

Finally, a 400 ns production run was performed under the NPT ensemble with random initial velocities, and trajectories were saved every 2 ps. Each system was simulated in three independent trials.

### 2.4 Identification of Binding Site Residues

Binding site residues were identified based on the activity ratio, defined as *n/N*, where *n* is the number of frames in which the minimum distance between any heavy atom of an amino acid residue and any heavy atom of the ligand is less than 5 Å, and *N* is the total number of frames. Residues with an activity ratio greater than 0.5 during the first 100 ns of simulation were considered part of the binding site, and the union of such residues across all ligand complexes was extracted (see Supplementary Figure S5). The RMSD trajectories for binding site residues in each production run are provided in Supplementary Figures S6, S7, and S8.

### 2.5 Trajectory Distance Calculation

For each protein or protein–ligand complex, the trajectories of the binding site residues were used to estimate probability density functions via kernel density estimation using a Gaussian kernel. The bandwidth was determined using Silverman’s rule-of-thumb. The similarity between probability density functions from different ligand systems was then evaluated using the Jensen–Shannon (JS) divergence. For systems *i* and *j*, the distance *W*_*ij*_ is defined as

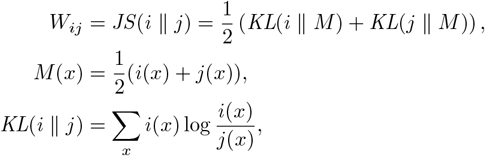

where *KL*(· ∥ ·) denotes the Kullback–Leibler divergence.

### 2.6 Binding Affinity Evaluation Based on Dynamic Similarity

The distance matrix *W* = {*W*_*ij*_} was embedded into a low-dimensional vector space. A set of 5-dimensional vectors was determined by minimizing the error function

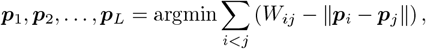

using simulated annealing and quasi-Newton methods. PCA was then applied to the embedded vectors, and the correlation between the first principal component (PC1) and the binding free energy Δ*G* was evaluated.

### 2.7 Estimation of the Sign of the PC1–Δ*G* Correlation

In the previous method [10], the correct orientation of PC1 with respect to Δ*G* was ambiguous without knowledge of the oracle Δ*G*. In our approach, a pseudo-oracle is obtained via docking simulations using AutoDock Vina [16]. For each protein–ligand pair, the docking score is calculated and its correlation with PC1 is determined. If PC1 is positively correlated with the docking score, we assume a positive correlation with Δ*G*, and vice versa. Although the docking score does not precisely match the experimental Δ*G*, it is sufficiently reliable to determine the polarity of the correlation.

### 2.8 Computational Environment

MD simulations were performed on the TSUBAME 4.0 GPU node (gpu 1 queue: AMD EPYC 9654 2.4 GHz [8 cores], NVIDIA H100 SXM5 94GB HBM2e [1 card], 96 GiB DDR5-4800 RAM) at the Center for Information Infrastructure, Institute of Science Tokyo. Amber22 [15] was used for MD simulations, and subsequent analyses were conducted in Python. The software libraries and versions used are listed below:

|  |  |
| --- | --- |
| Amber | 22 Update5 |
| AmberTools | 23 Update6 |
| AutoDock Vina | 1.2.5 |
| Python | 3.9.19 |
| NumPy | 1.26.4 |
| MDAnalysis | 2.7.0 |
| scikit-learn | 1.3.0 |
| SciPy | 1.12.0 |

## 3 Results and Discussion

### 3.1 Comparison of Complex Dynamics Using JS Divergence

Figure 2 shows the distance matrix computed from the JS divergence between the probability distributions derived from the binding site trajectories. It is evident that the distances between the apo system (system 0) and the ligand-bound systems are larger compared to those among the complexes, indicating significant dynamic differences induced by ligand binding.

**Figure 2:**
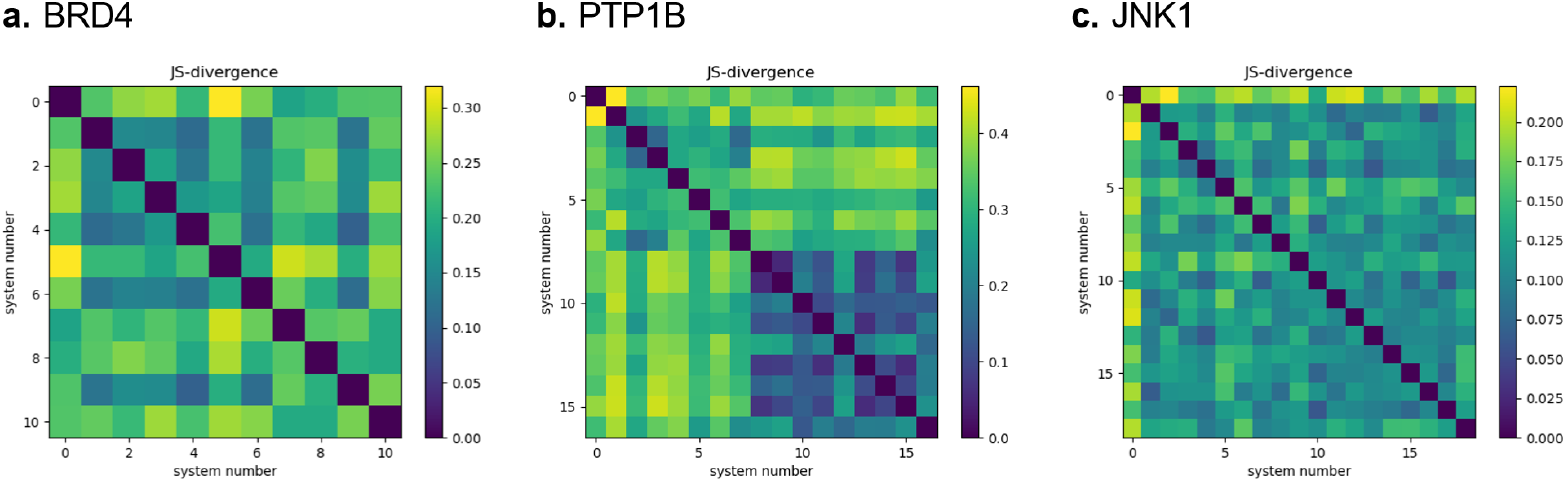
Distance matrix based on the JS divergence between probability distributions from the binding site trajectories. Higher (yellow) values indicate greater dynamic differences. System number 0 represents the apo protein trajectory.

### 3.2 Principal Component Analysis of Embedded Vectors

The distance matrix *W* was embedded into a 5-dimensional vector space and subjected to PCA. As shown in Figure 3, for all datasets the ligand systems with high and low Δ*G* values are clearly separated along PC1.

**Figure 3:**
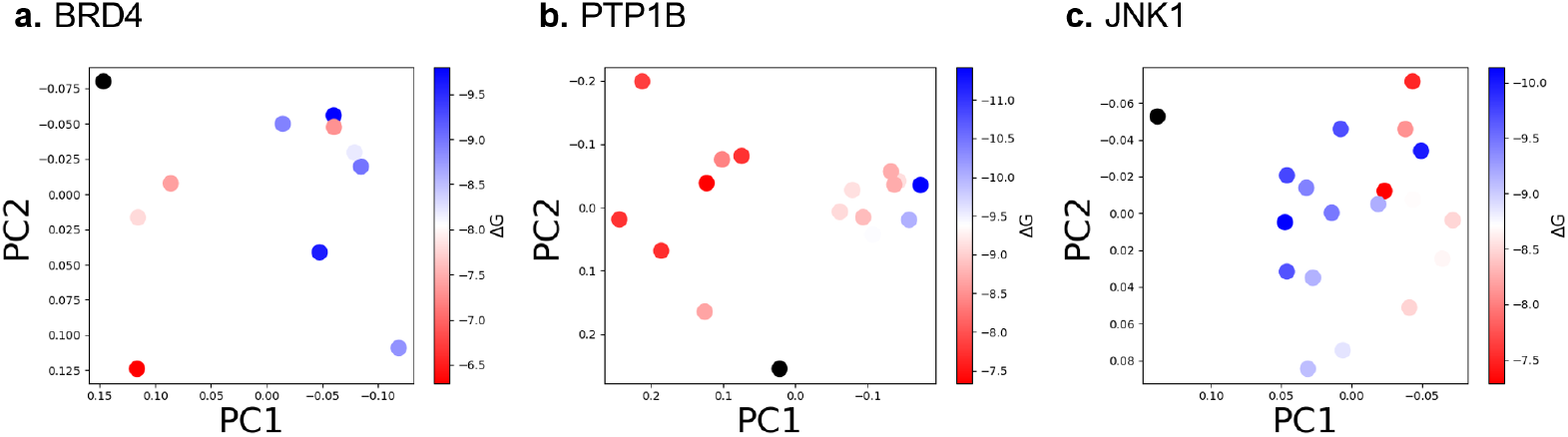
PCA results after embedding the distance matrix *W* into a 5-dimensional space.

### 3.3 Binding Affinity Evaluation Based on Dynamic Similarity

The correlation between PC1 and the oracle Δ*G* is presented in the upper panel of Figure 4. For the BRD4 and PTP1B datasets, PC1 exhibits a positive correlation with Δ*G*, whereas for the JNK1 dataset, a negative correlation is observed.

**Figure 4:**
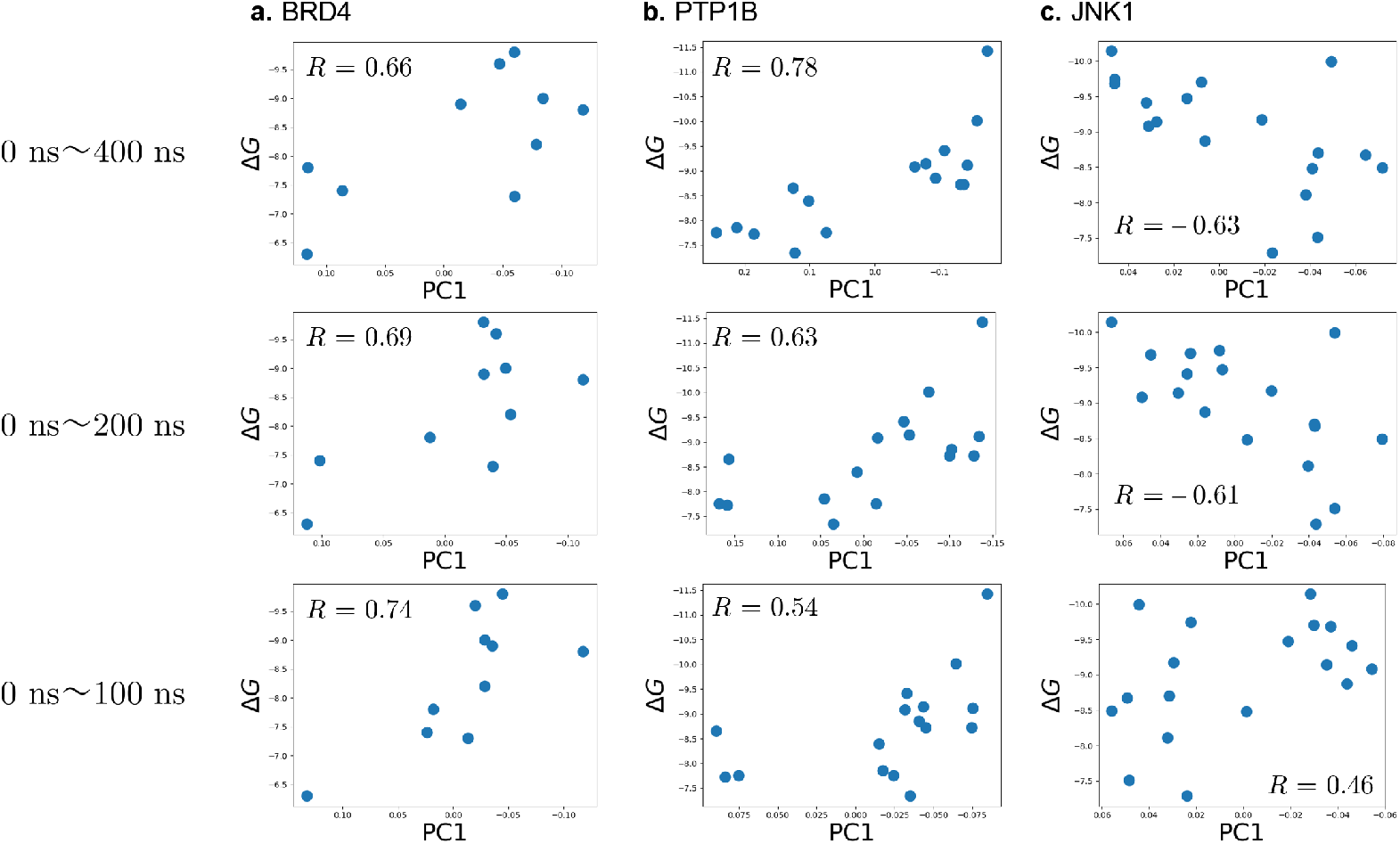
Correlation plots of PC1 with the oracle Δ*G*. The upper panel shows results using trajectories up to 400 ns, while the middle and lower panels correspond to shorter simulation durations.

### 3.4 Effect of MD Simulation Length on Prediction Accuracy

We investigated the impact of production run simulation time by comparing trajectories of 0–400 ns, 0–200 ns, and 0–100 ns (Figure 4). For the BRD4 dataset, the separation of ligand systems by Δ*G* along PC1 is evident in all cases. However, for PTP1B and JNK1, while 200 ns and 400 ns trajectories yield clear separation between high and low Δ*G* systems, the 100 ns trajectories do not. This suggests that a simulation time of around 200 ns or more is necessary to capture the dynamic behavior reflecting structural stabilization and binding affinity differences.

### 3.5 Estimation of the PC1–Δ*G* Correlation Sign

Figure 5 (upper panel) compares PC1 with the docking scores Δ*G*_dock_ obtained via AutoDock Vina. For BRD4 and PTP1B, the signs of the correlation coefficients between PC1 and Δ*G*_dock_ match those between PC1 and the oracle Δ*G*, validating the use of docking scores for sign determination. For JNK1, however, an inverse sign was observed. The lower panel of Figure 5 shows that although the docking scores do not perfectly agree with the oracle Δ*G*, they are sufficiently reliable for determining the polarity of the PC1–Δ*G* correlation. In cases where a few compounds have been evaluated using absolute binding free energy calculations such as FEP or MP-CAFEE [17], those values could also be used to determine the sign.

**Figure 5:**
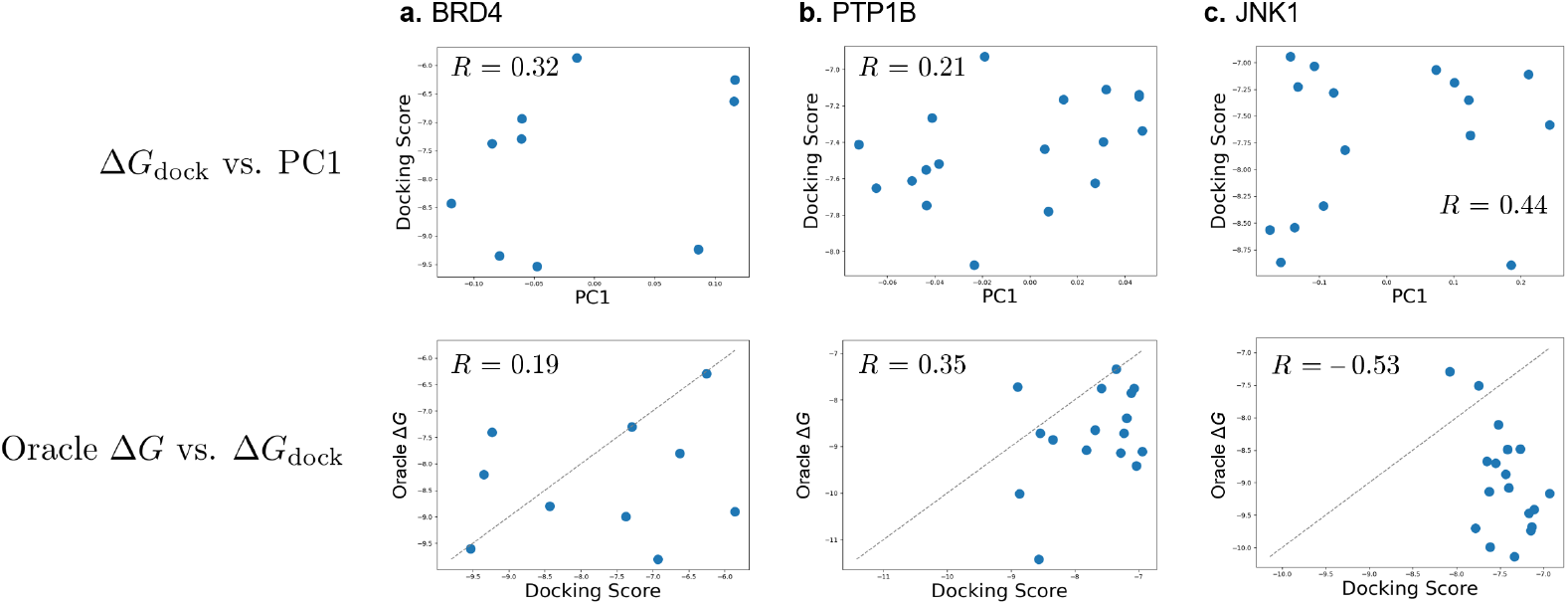
(Upper panel) Relationship between PC1 and docking scores Δ*G*_dock_. (Lower panel) Comparison between docking scores Δ*G*_dock_ and oracle Δ*G*.

### 3.6 Comparison of Prediction Accuracy and Computational Cost

Table 1 compares the computational cost and performance between the method of Yasuda et al. [10] and the present work. In the previous approach, deep neural network-based Wasserstein distance calculations required several hours per ligand pair using an NVIDIA GeForce RTX 3090 24GB GPU, whereas our method employing JS divergence computes the distance in a matter of seconds. Although the correlation between PC1 and Δ*G* is comparable, our method reduces both the required simulation time (200 ns instead of 400 ns) and the per-comparison computational cost.

**Table 1:** Comparison with the previous study [10].

|  | BRD4 |  | PTP1B |  |
| --- | --- | --- | --- | --- |
|  | Previous Method | This Study | Previous Method | This Study |
| Correlation (PC1 vs. $\Delta G$) | $R = 0.88$ | $R = 0.66$ | $R = 0.70$ | $R = 0.78$ |
| Required Simulation Length | 400 ns | 200 ns | 400 ns | 200 ns |
| Similarity Computation | DNN-based Wasserstein distance | JS divergence | DNN-based Wasserstein distance | JS divergence |
| Time per Pair Comparison | Several hours | Several seconds | Several hours | Several seconds |

Additionally, we compared our method with relative FEP calculations for the PTP1B and JNK1 datasets [18]. Although direct comparison is challenging due to differences in simulation protocols (FEP simulations for PTP1B and JNK1 required total simulation times of approximately 3000 ns and 2300 ns, respectively, compared to around 6800 ns and 7600 ns in our study), the correlations (*R*^2^ values) were comparable (FEP: *R*^2^ = 0.47 for PTP1B and *R*^2^ = 0.74 for JNK1; this study: *R*^2^ = 0.61 for PTP1B and *R*^2^ = 0.48 for JNK1). It is important to note that while relative FEP directly computes free energy differences (ΔΔ*G*), our method only ranks the systems by Δ*G*. Nevertheless, for targets like BRD4 where ligand structures are highly diverse and relative FEP may struggle, the proposed method offers a viable alternative.

## 4 Conclusion

In this study, we have presented improvements to an MD simulation-based method for protein–ligand binding affinity prediction. By introducing the Jensen–Shannon divergence, we can rapidly compute the distance matrix between system trajectories without the need for deep learning, thereby significantly reducing computational cost. We also demonstrated that the production run simulation time can be reduced from 400 ns to 200 ns while maintaining comparable accuracy. Finally, by incorporating coarse Δ*G* estimations from AutoDock Vina, we proposed a strategy to determine the sign of the PC1–Δ*G* correlation.

Although our method requires three-dimensional structural information of the target, the increasing accuracy of structure prediction tools such as AlphaFold3 [19] suggests that our approach may soon be applicable even to targets with unresolved experimental structures. Future work will explore the application of our method using predicted structures.

## Author Contributions

K.I. was responsible for executing the research, implementing and analyzing the work, visualizing the results, drafting the article, and revising the manuscript. M.O. provided overall research supervision, conceived and designed the work, led the discussion of the results, contributed to drafting the article, and revised the manuscript. All authors approved the final version to be published.

## Acknowledgements

This work was supported by the JST FOREST (JPMJFR216J) and AMED BINDS (JP24ama121026). The authors thank Ikki Yasuda for valuable discussions regarding the computational cost of the deep neural network-based Wasserstein distance calculation, and Kairi Furui for contributions to the discussion of computational time in FEP calculations.

