## Supporting Information for "Protein–Ligand Affinity Prediction via Jensen–Shannon Divergence of Molecular Dynamics Simulation Trajectories"

**a.** BRD4 **b.** PTP1B **c.** JNK1

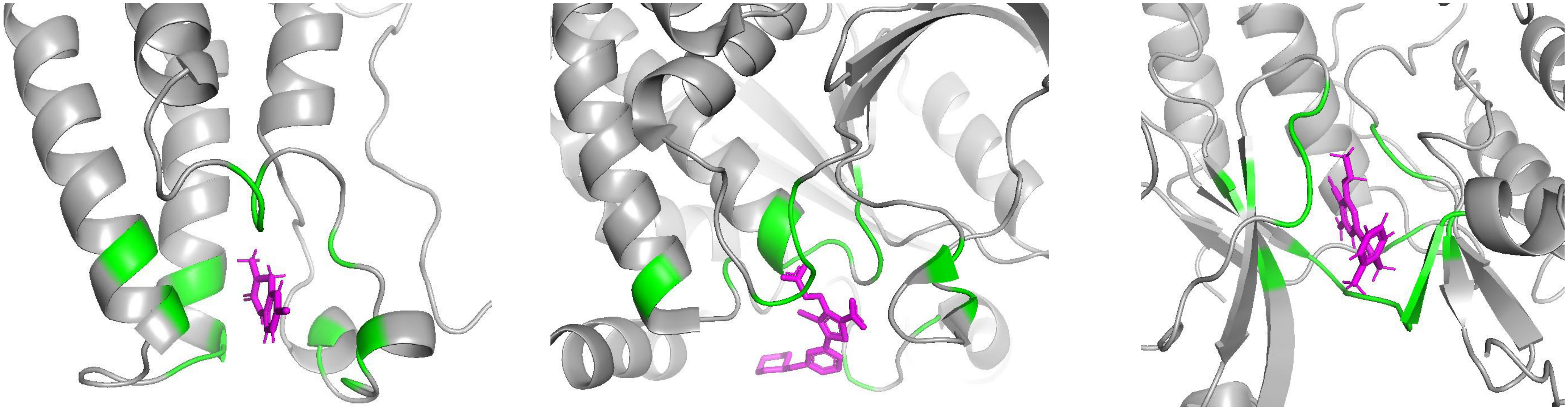

**Supplementary Figure S1**. Complex structures for each target

### bromodomain 4 (BRD4)

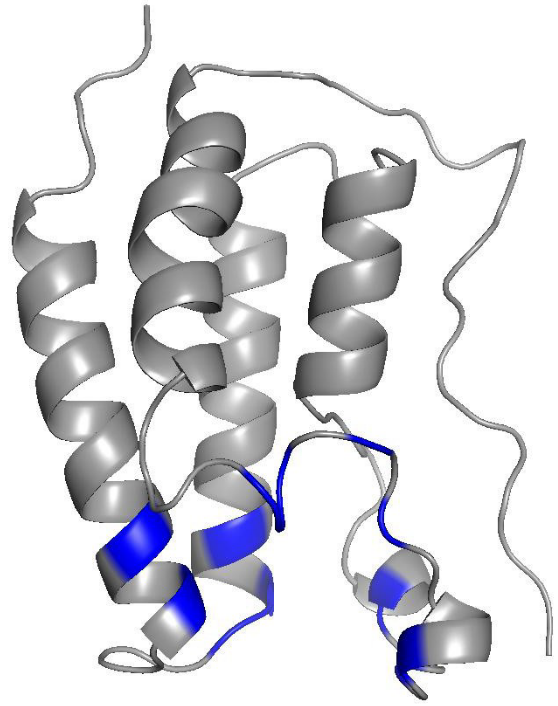

PDB ID: 2OSS

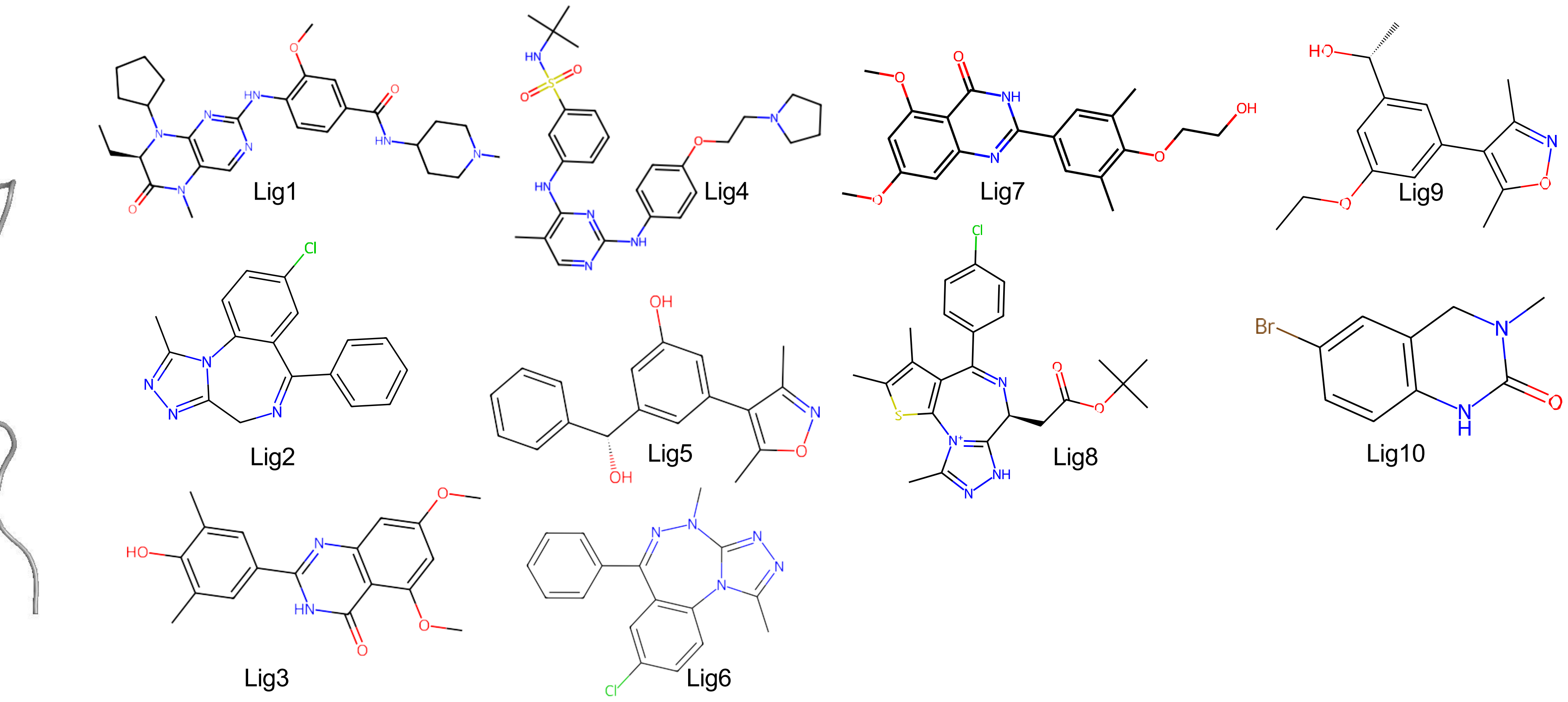


**Supplementary Figure S2**. BRD4 protein three-dimensional structure (PDB) and the structure of the compounds used.

### protein tyrosine phosphatase 1B (PTP1B)

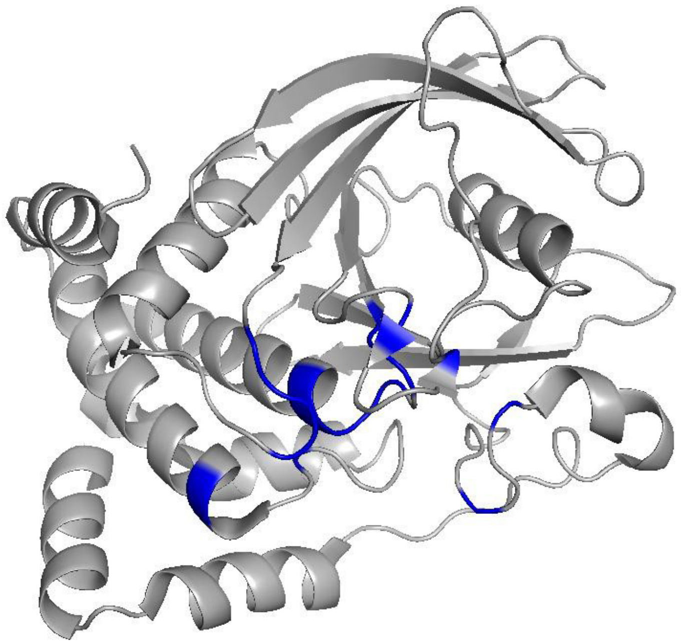

PDB ID: 2QBS

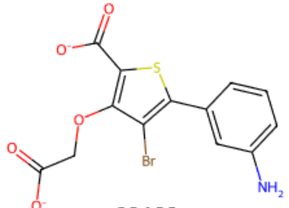

23466

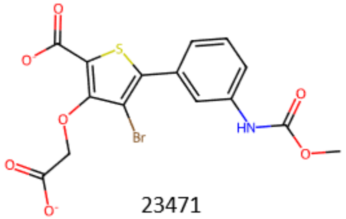

23471

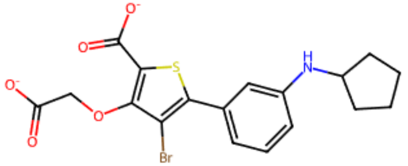

23475

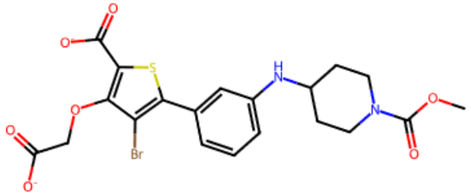

23480

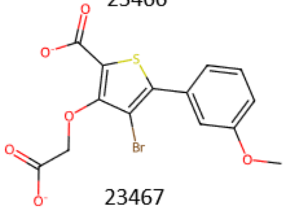

23467

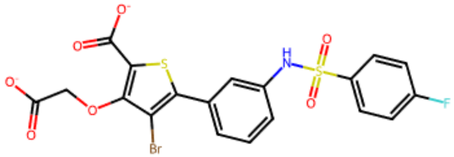

23472

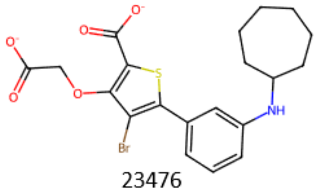

23476

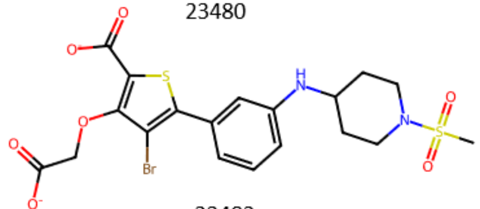

23482

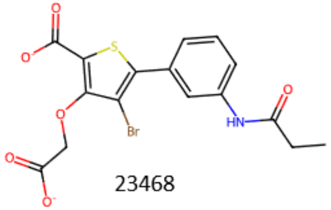

23468

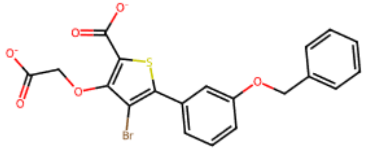

23473

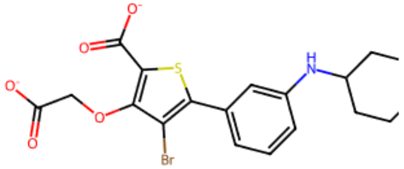

23477

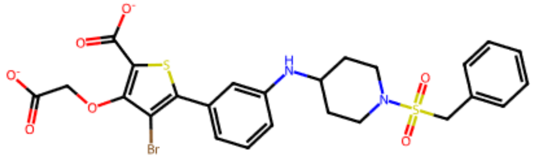

20667

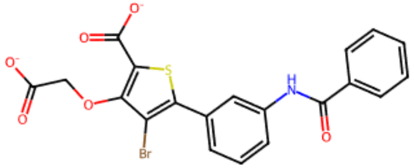

23469

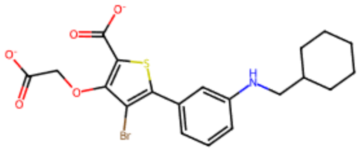

23474

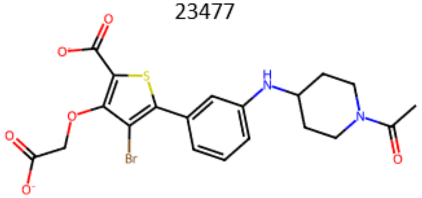

23479

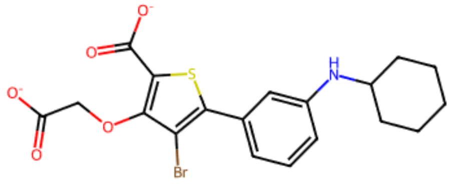

20670

**Supplementary Figure S3**. PTP1B protein three-dimensional structure (PDB) and the structure of the compounds used.

### c-Jun N-terminal kinase 1 (JNK1)

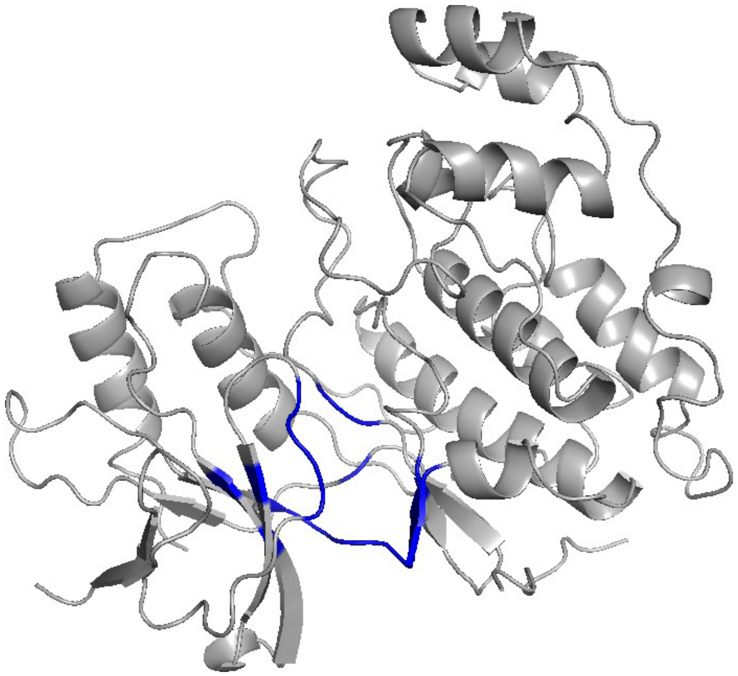

PDB ID: 2GMX

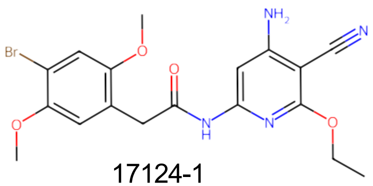

17124-1

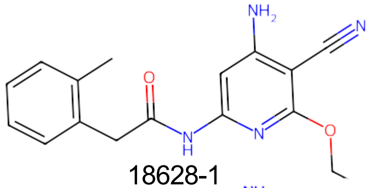

18628-1

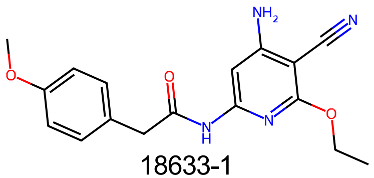

18633-1

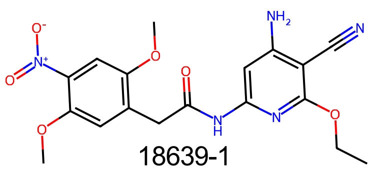

18639-1

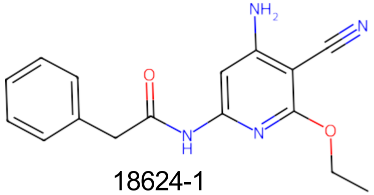

18624-1

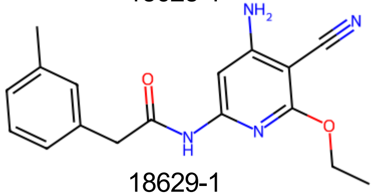

18629-1

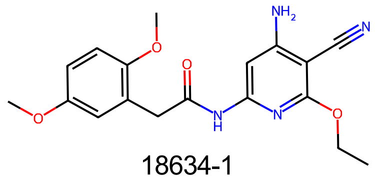

18634-1

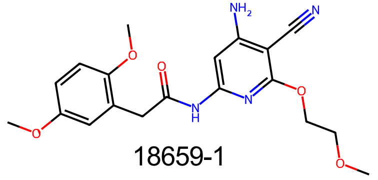

18659-1

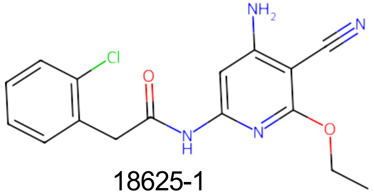

18625-1

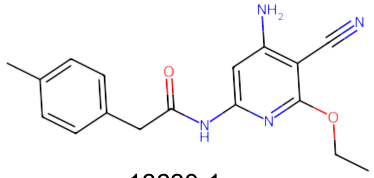

18630-1

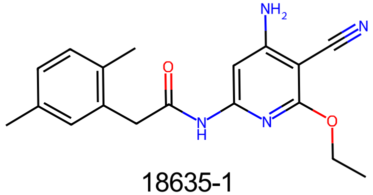

18635-1

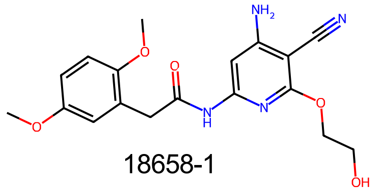

18658-1

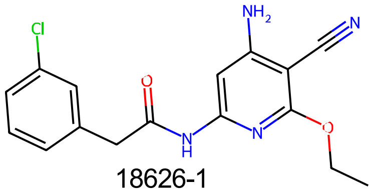

18626-1

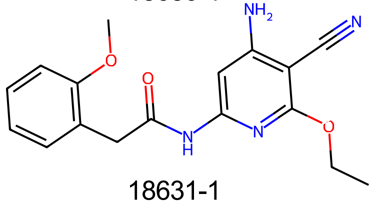

18631-1

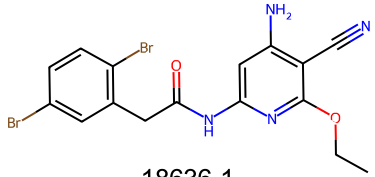

18636-1

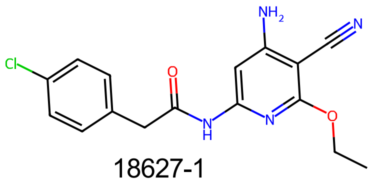

18627-1

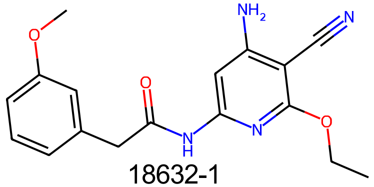

18632-1

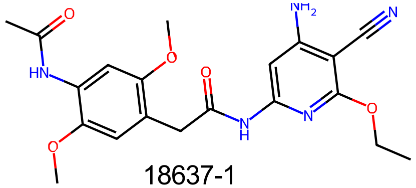

18637-1

**Supplementary Figure S4**. JNK1 protein three-dimensional structure (PDB) and the structure of the compounds used.

**a.** BRD4 **b.** PTP1B **c.** JNK1

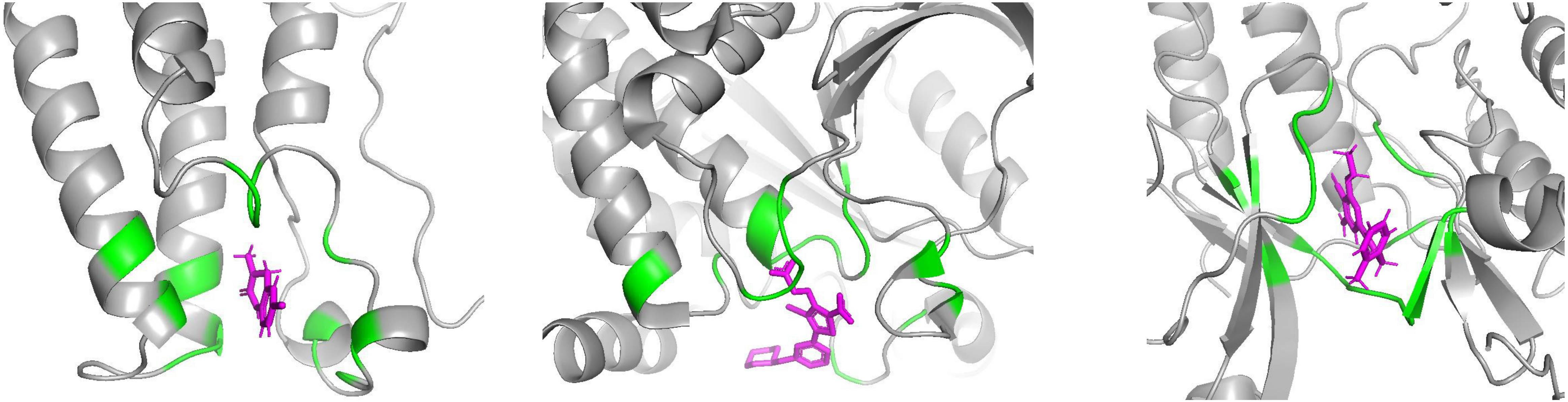

**a. BRD4 — 13 residues**

| | |
|---|---|
| Trp81 | Tyr97 |
| Pro82 | Cys136 |
| Phe83 | Tyr139 |
| Gln85 | Asn140 |
| Val87 | Ile146 |
| Leu92 | Met149 |
| Leu94 | |

**b. PTP1B — 17 residues**

| | |
|---|---|
| Tyr45 | Ser215 |
| Val48 | Ala216 |
| Glu114 | Ile218 |
| Lys119 | Gly219 |
| Pro179 | Arg220 |
| Asp180 | Ser221 |
| Phe181 | Gln261 |
| Gly182 | Gln265 |

**c. JNK1 — 18 residues**

| | |
|---|---|
| Ile25 | Gln102 |
| Gly26 | Leu103 |
| Ser27 | Met104 |
| Gly28 | Asp105 |
| Val33 | Ala106 |
| Ala46 | Asn107 |
| Lys48 | Val151 |
| Ile79 | Leu161 |
| Met101 | Asp162 |

**Supplementary Figure S5**. Amino acid residues identified as binding site residues. The green regions in the three-dimensional ribbon diagram indicate the binding site.

### RMSD of binding site residues (3 trials), BRD4

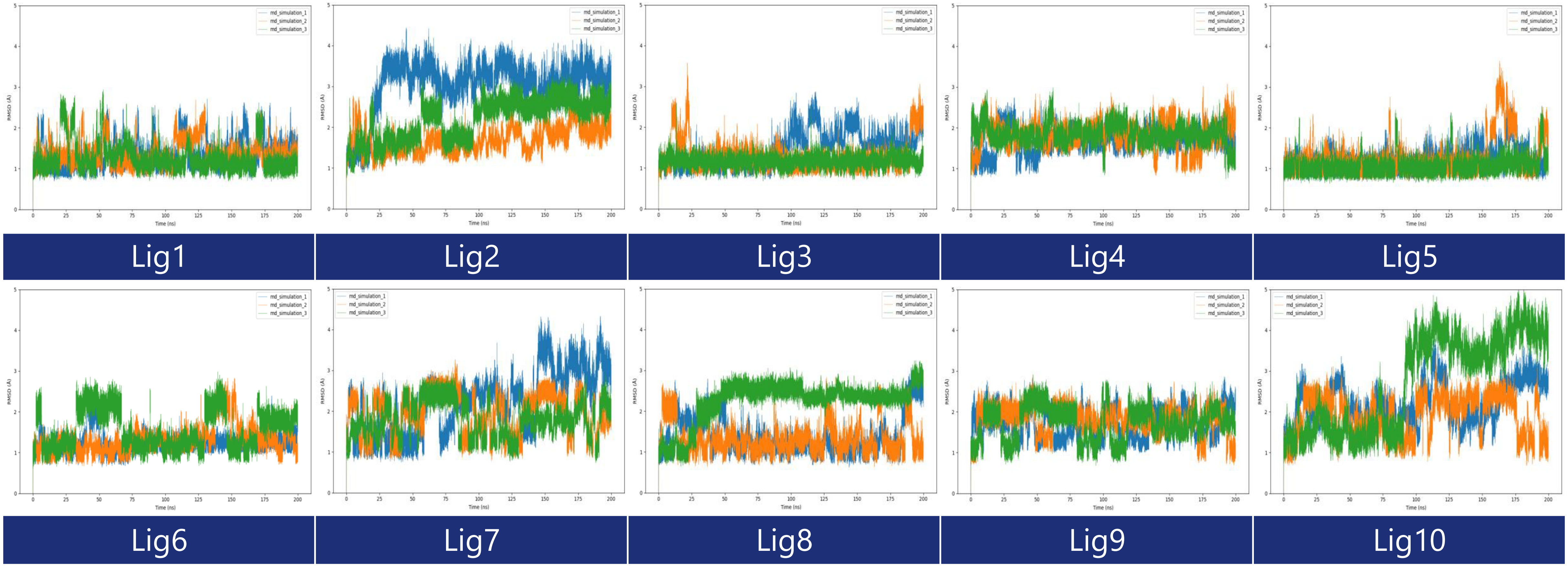


**Supplementary Figure S6**. RMSD of binding site residues for each BRD4 protein system (3 MD trials)

### RMSD of binding site residues (3 trials), PTP1B

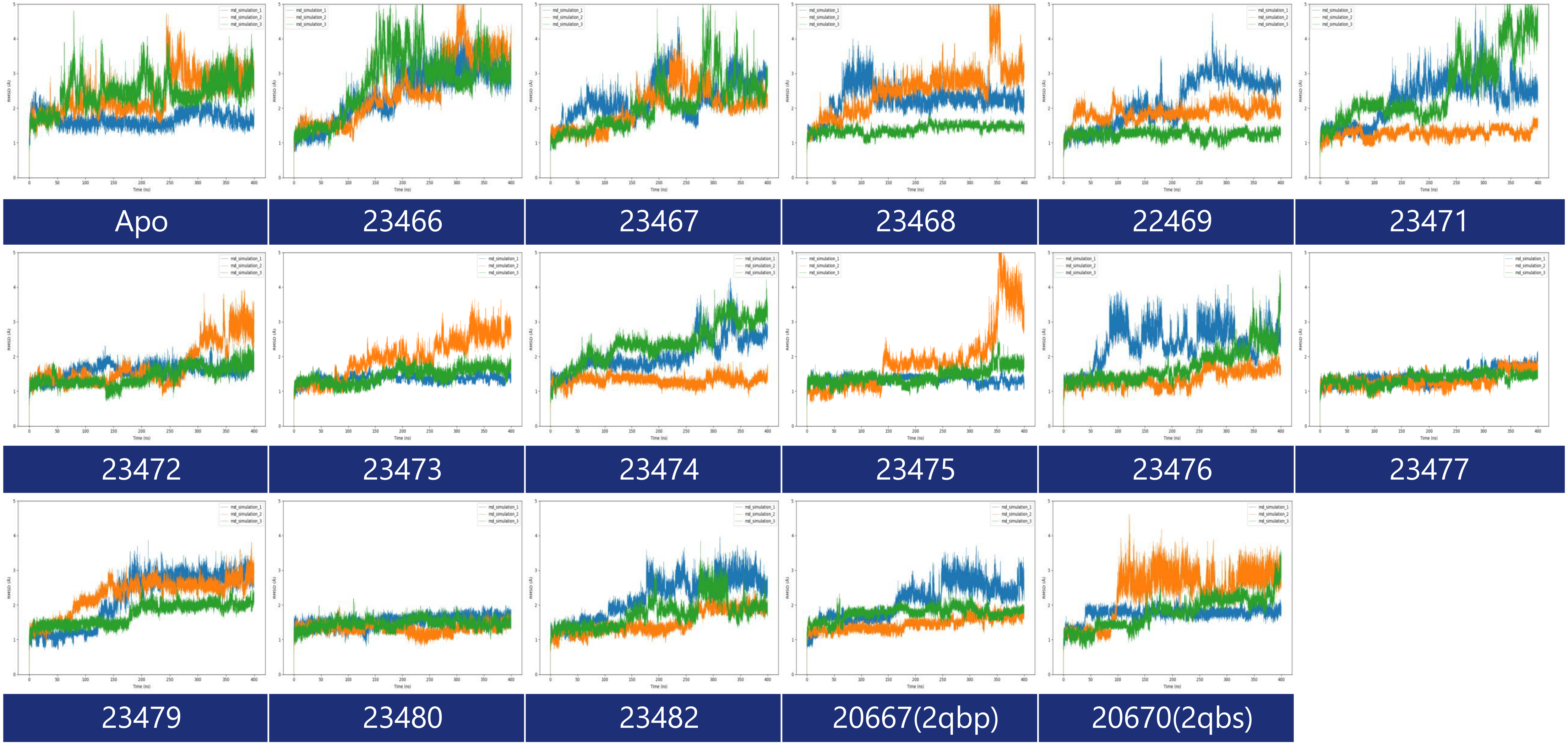


**Supplementary Figure S7**. RMSD of binding site residues for each PTP1B protein system (3 MD trials)

#### RMSD of binding site residues (3 trials), JNK1

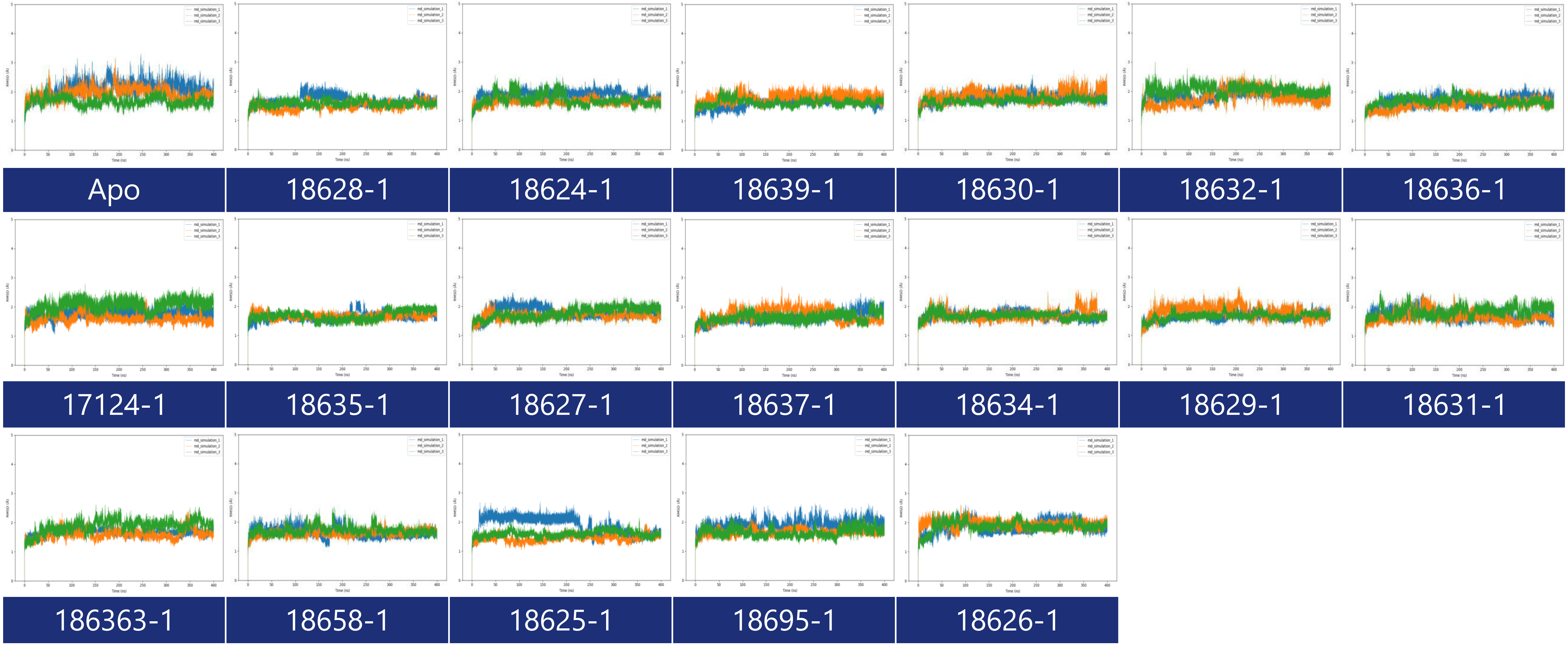


**Supplementary Figure S8**. RMSD of binding site residues for each JNK1 protein system (3 MD trials)
